# mTOR-driven widespread exon skipping renders multifaceted gene regulation and proteome complexity

**DOI:** 10.1101/2020.02.27.967737

**Authors:** Sze Cheng, Naima Ahmed Fahmi, Meeyeon Park, Jae-Woong Chang, Jiao Sun, Kaitlyn Thao, Hsin-Sung Yeh, Wei Zhang, Jeongsik Yong

## Abstract

The mammalian target of rapamycin (mTOR) pathway is crucial in cell proliferation. Previously, we reported transcriptome-wide 3’-untranslated region (UTR) shortening by alternative polyadenylation upon mTOR activation and its impact on the proteome. Here, we further interrogated the mTOR-activated transcriptome and found that hyperactivation of mTOR promotes transcriptome-wide exon-skipping/exclusion, producing short isoform transcripts from genes. This widespread exon skipping confers multifarious regulations in the mTOR-controlled functional proteomics: alternative splicing (AS) in the 5’-UTR controls translation efficiency while AS in coding regions widely affects the protein length and functional domains. They also alter the half-life of proteins and affect the regulatory post-translational modifications. Among the RNA processing factors differentially regulated by mTOR signaling, we found that SRSF3 mechanistically facilitates exon skipping in the mTOR-activated transcriptome. This study reveals a role of mTOR in AS regulation and demonstrates that widespread AS is a multifaceted modulator of the mTOR-regulated functional proteome.

## Introduction

mRNA splicing is a critical co-transcriptional process to produce uninterrupted coding DNA sequences (CDS) for protein translation^1^. Alternative splicing (AS) can occur by alternating the inclusion of exons or part of exons to produce mature mRNAs^2–4^. Thus, AS diversifies the proteome by potentiating the production of multiple protein isoforms from a gene^5,6^. Although AS serves as an essential layer of post-transcriptional regulation of gene expression in eukaryotes^7,8^, much remains to be explored concerning its regulation and impact on the resulting proteome.

Mammalian target of rapamycin (mTOR) is a serine/threonine kinase and is the major component of the mTOR Complex 1 (mTORC1) and 2 (mTORC2)^9–11^. Heterodimeric tuberous sclerosis complex composed of TSC1 and TSC2 is a negative regulator of mTORC1. mTOR promotes the translation of a subset of mRNAs that contain the 5’-terminal oligopyrimidine tract (5’-TOP)^12–15^. mTOR was also shown to regulate select RNA-binding proteins (RBPs) and functions in post-transcriptional regulation^16–18^. Hyperactivation of mTOR leads to transcriptome-wide alternative polyadenylation (APA) in the 3’-untranslated regions (3’-UTR) of transcripts and affects diverse cellular pathways^19,20^. mTOR activation also prefers the expression of splicing factor U2AF1a isoform to that of the b isoform and renders U2AF1a-dependent AS events^21^.

In this study, we investigated mTOR-driven transcriptome changes in alternative splicing and their impact on the resulting proteome using various cellular models. We found that widespread exon skipping by mTOR activation diversifies the proteome by changing functional domains, post-translational modifications, and protein stabilities.

## Results and Discussion

### mTOR activation leads to transcriptome-wide exon skipping

To better understand mTOR-driven transcriptomic features, we profiled the transcriptome of various mammalian cells with low and high mTOR contents: *Tsc1*^*-/-*^ mouse embryonic fibroblasts (MEFs), MCF7 and MDA-MB-361 cells treated with DMSO (mock) or Torin 1, a potent inhibitor of mTOR. From our initial interrogation that focused on the RNA processing pathway, we found numerous RBPs involved in AS and APA to be differentially expressed, suggesting the comprehensive role of mTOR in the transcriptome (Fig. 1A and Suppl Table 1). We then profiled AS events using the custom-developed AS-Quant pipeline^22^. Strikingly, we found that chemical inhibition of mTOR in *Tsc1*^-/-^ MEFs drives exon inclusion in general: among the affected AS events, exon inclusion in cassette type AS was almost exclusively preferred (497 events; 96%) in Torin 1-treated *Tsc1*^-/-^ MEFs (Fig. 1B and Suppl Table 2). This trend of exon inclusion upon mTOR inhibition was consistent with other types of AS events with varying degrees (59% in 5’-AS site and 82% in 3’-AS site) (Fig. 1B, Suppl Table 2). Consistent with these findings, comparison of other pairs of AS using the datasets from WT, *Tsc1*^-/-^ MEFs, and human breast cancer cell lines MCF7 and MDA-MB-361 showed an overall increase of exon inclusion upon mTOR inhibition. (Fig. 1C, Suppl Fig. 1A-B, and Suppl Table 3-4). We noticed that the number of exon inclusions varies among profiled cellular models (60% in breast cancer transcriptome compared to 90% in *Tsc1*^-/-^ MEF transcriptome). This discrepancy may come from the degree of mTOR activation in these cell lines examined: *Tsc1*^-/-^ MEFs are genetically programmed to hyperactivate mTOR signaling while MCF7 and MDA-MB-361 cells may have a reduced level of mTOR hyperactivation. Thus, upon mTOR inhibition, changes in AS events are maximized in the *Tsc1*^-/-^ system compared to breast cancer cells. In a similar context, when the same *Tsc1*^-/-^ transcriptome was analyzed against the WT (basal mTOR activity) transcriptome, it showed less exon inclusion-biased AS events compared to that of the *Tsc1*^-/-^ transcriptome with Torin 1 treatment. We further delved into the nucleotide contexts surrounding the 3’-splice site and found that mTOR activation prefers a longer stretch of poly-pyrimidine tract (−3 to -20 position) in the cassette exon and a weak poly-pyrimidine tract with biased C at the -1 position in the alternative 3’-splice site in both *Tsc1*^-/-^ MEFs and human cancer cells (Suppl Fig. 1C). Further examinations of affected AS events showed that the majority of them occur in the CDS of mRNA (65-68%) while 26-27% of them are found in the 5’-UTR (Fig. 1D and Suppl Fig. 1D). These findings suggest that mTOR activation escalates the frequency of exon skipping during pre-mRNA splicing leading to preferential expression of short isoforms in the transcriptome.

**Figure 1.**
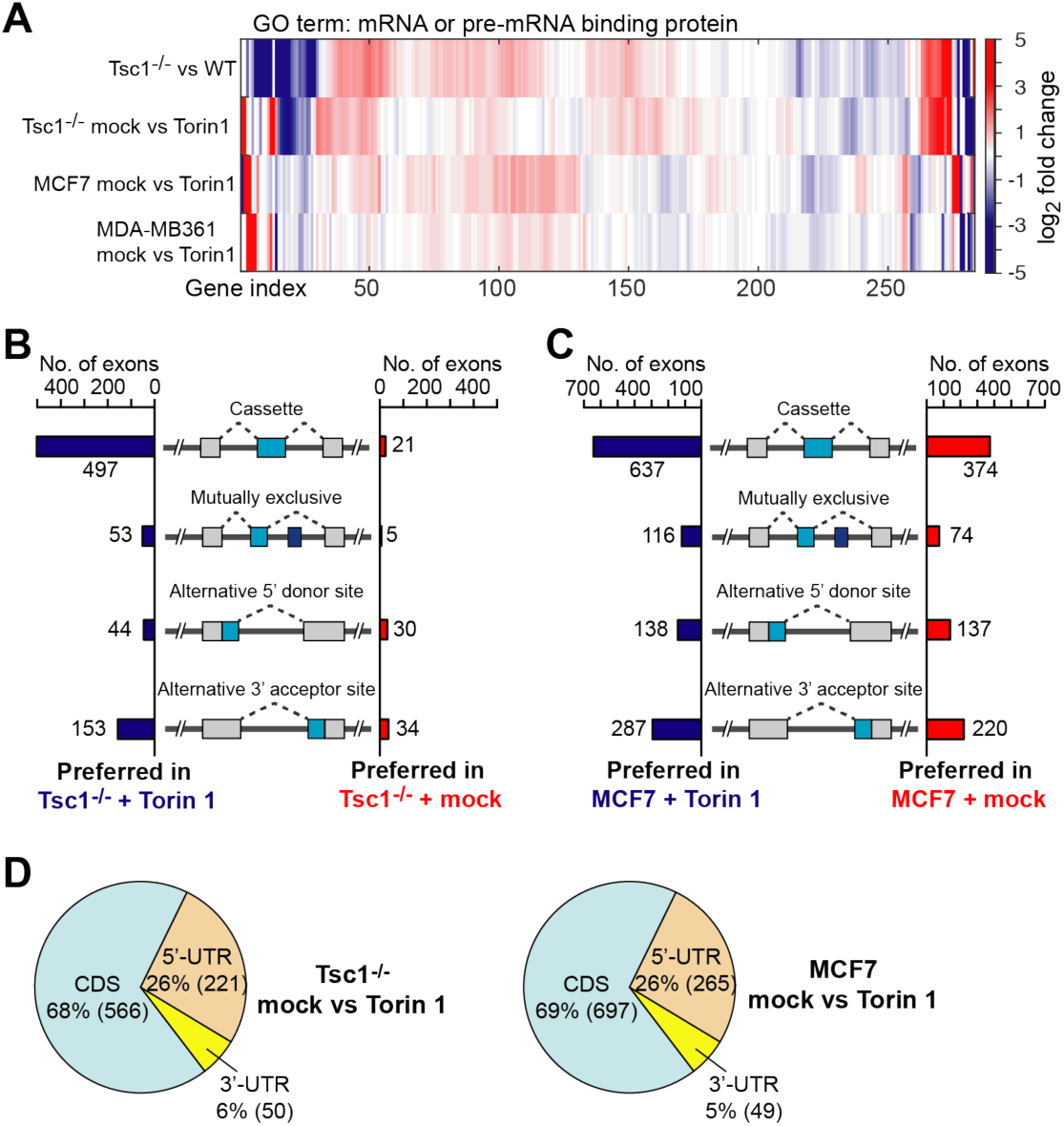
The mTOR-activated transcriptome features widespread exon skipping. A) Hierarchical clustering of select RNA-binding proteins (RBPs) using differential gene expression analysis in low and high mTOR cellular environments. Comparisons were made in the combination of cells as indicated. The x-axis indicates the 285 different RBPs analyzed. The y-axis shows the combination of comparisons. The expression level is color-coded as shown in the scale bar and represents the log2 fold changes of gene expression. B-C) Analyses of four different types of alternative splicing events in B) *Tsc1*^-/-^ mock vs. Torin 1 treatment and C) Breast cancer cell line MCF7 mock vs. Torin 1 treatment. Cases of alternative exon inclusion are compared and illustrated in the combination as indicated in the figure. D) Distribution of mTOR-regulated alternative splicing (AS) events in different regions of the mRNA.

### mTOR regulates translation through 5’-UTR AS

Next, we validated mTOR-regulated AS events on select genes using PCR and analytical gel electrophoresis. Consistently, all tested genes showed exon skipping upon mTOR activation (*Tsc1*^-/-^ MEFs mock vs Torin 1 treatment and *Tsc1*^-/-^ vs WT MEFs) (Fig. 2A, Suppl Fig. 2A). Interestingly, we found that a number of AS events in CDS where the exon-inclusion transcript contains a premature termination codon (PTC). Conversely, we also found cases where exon-inclusion converts a PTC-containing isoform to an isoform with a full open reading frame. (Fig. 2B, Suppl Fig. 2B and C). In *Mdm4*, whose expression is important for the stability of p53^23^, mTOR activation leads to the expression of evolutionary conserved PTC-containing *Mdm4* transcript (NR_126506) by skipping exon 6 while mTOR inhibition switches back to the expression of full open reading frame (ORF) (NM_001302803) (Fig. 2B). Importantly, the PTC-containing *Mdm4* transcript degrades faster than the coding transcript (Fig. 2C and Suppl Fig. 2D). This suggests that mTOR-driven AS switches on and off gene expression by introducing PTC in select genes (Suppl Fig. 2B).

**Figure 2.**
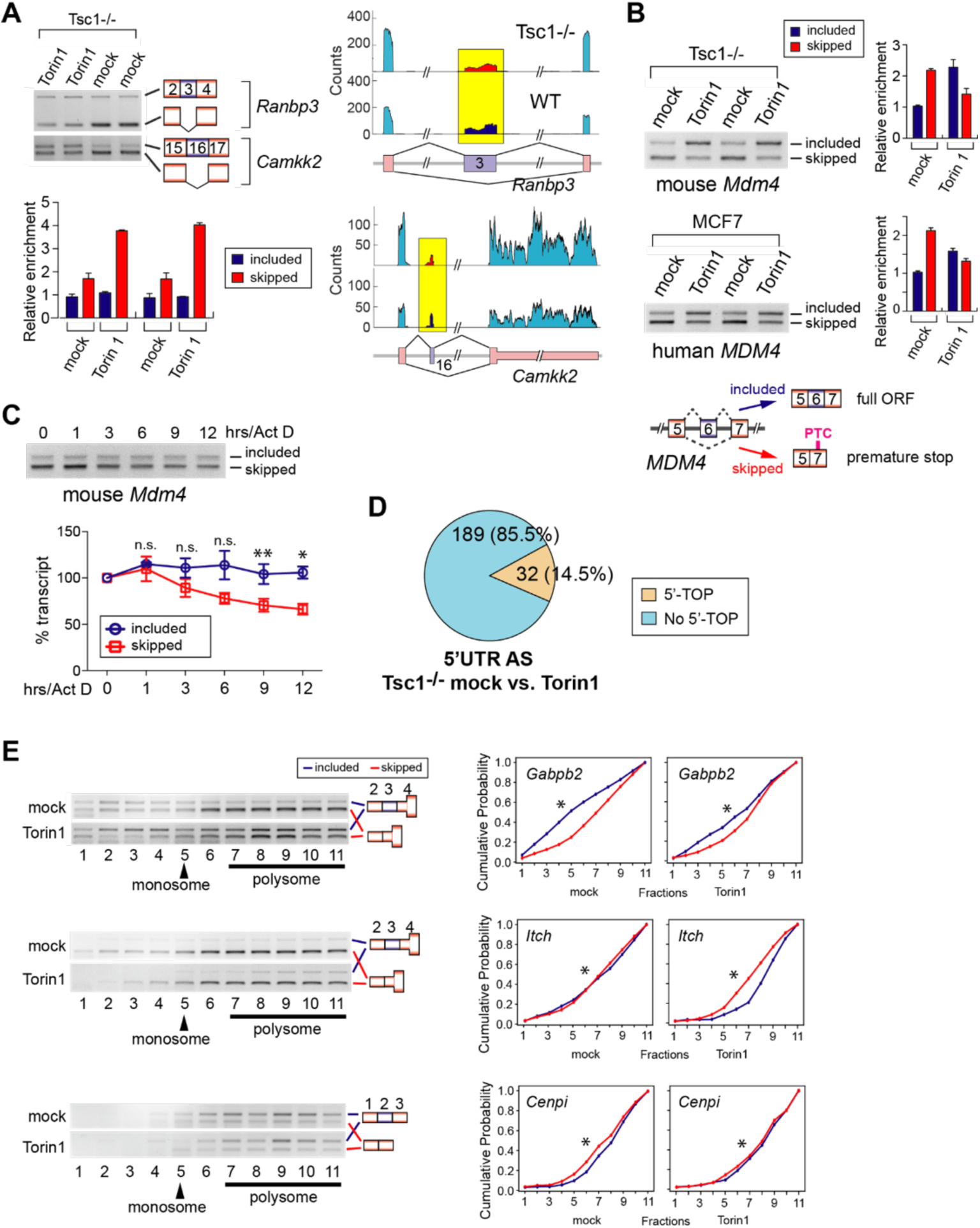
Exon skipping highlights a regulatory role of non-coding regions in the mTOR-activated proteome. A) Semi-quantitative RT-PCR validation of mTOR-regulated AS events. (left panel) RT-PCR and analytical gel electrophoresis were conducted in mock- and 50 nM Torin 1-treated *Tsc1*^-/-^ MEFs. Two technical repeats were done. Quantitation of gel images was done by densitometry in ImageStudioLite software. (right panel) RNA-Seq read alignments of *Ranbp3* and *Camkk2* AS events in mock- and Torin 1-treated *Tsc1*^-/-^ MEFs. Affected AS exon is highlighted in yellow. B) Semi-quantitative RT-PCR validation of full open reading frame (ORF) containing and premature termination codon (PTC)-containing transcripts. *Mdm4* AS events were analyzed in *Tsc1*^-/-^ MEFs in the presence or absence of Torin 1 treatment (50 nM). The same analysis was also conducted for human *MDM4* AS using MCF7 cells. The relative quantification of the full ORF- and PTC-containing transcripts was done by densitometry in ImageStudioLite software. Two technical repeats were done. C) Stability of *Mdm4* isoform transcripts. Semi-quantitative RT-PCR analysis was performed using *Tsc1*^-/-^ MEFs treated with 1 ug/ml actinomycin D for indicated time points. Quantitation was done by densitometry in ImageStudioLite software. The data are represented as mean (SD). Statistical analysis was done using two technical replicates. *p=0.0232, **p=0.0657. n.s. denotes no significance. D) Proportion of 5’-UTR AS events containing the 5’-TOP motif. E) Polysome analyses of mTOR-regulated 5’-UTR AS transcripts *Gabpb2* (exon-included: NM_029885, exon-skipped: NM_172512), *Itch* (exon-included: NM_001243712, exon-skipped: NM_008395), and *Cenpi* (exon-included: NM_145924, exon-skipped: NM_001305631). Semi-quantitative RT-PCR and analytical gel electrophoresis were performed on the cytoplasmic fractions of mock- and Torin 1-treated *Tsc1*^-/-^ MEFs. The monosome and polysome fractions are indicated. A schematic of the AS event for each transcript is shown on the right of the gel images. The cumulative distribution of the AS transcripts in each fraction is plotted against the summation of that isoform in all 11 fractions. The x-axis indicates the fraction number and y-axis indicates the cumulative probability. For all comparisons, p<1.0e-10.

Besides the 5’-TOP feature of mRNAs^24,25^, the 5’-UTR length plays a role in mTOR-regulated translation^26,27^. Interrogation of 5’-UTR sequences found that the majority of genes showing mTOR-driven 5’-UTR AS do not contain the 5’-TOP motif (Fig. 2D). To examine the functional relevance of mTOR-modulated 5’-UTR length regulation in translation, two 5’-UTR AS isoforms lacking the 5’-TOP motif were analyzed in polysome profiling using analytical gel electrophoresis and cumulative plots (Fig. 2E and Suppl Fig. 2E-H). Cumulative plots of polysome profiling could determine statistically significant changes in the distribution of transcripts as exemplified by 5’-TOP containing transcripts *Eef2* and *Hsp90ab1* (Suppl Fig. 2G). All tested genes showed a difference in the polysome formation by the 5’-UTR AS: a short isoform of *Gabpb2* formed polysome more efficiently and sustained polysome formation more rigorously upon Torin 1 treatment compared to its longer isoform. In contrast, a long isoform of *Itch* and *Cenpi* showed the characteristics of *Gabpb2* short isoform in the polysome formation upon mock and Torin 1 treatment (Fig. 2E). The number of upstream open-reading-frames (uORFs) plays a critical role in determining the translation efficiency, and it has been shown to positively correlate with the length of the 5’-UTR^28,29^. However, our studies suggest that there is no unanimous theme of long vs. short in the formation or dissociation of polysome. This could be explained by exons being relatively short in length and AS may not be able to substantially modify the number of uORFs in the 5’-UTR. Overall, these results suggest that 5’-UTR AS is a previously unrecognized element in translational regulation by mTOR and propose that it could be a resistance mechanism to mTOR inhibitors.

### AS contributes to the mTOR-programmed functional proteome at multiple levels

To understand the functional relevance of mTOR-regulated AS in CDS, we first linked the affected exons to Pfam domains and used them for pathway analyses. From this we found 114 affected Pfam domains that are associated with various KEGG pathways and GO terms (Fig. 3A; Suppl Fig. 3A-B). AS of *Sirt2* and *Mdm2* in the CDS causes a frameshift in the original ORF and enforces the use of the second most available methionine for translation initiation. These events produce N-terminal truncated protein isoforms as illustrated in Fig. 3B and Suppl Fig. 3C. Cycloheximide chase assays on Flag-tagged SIRT2 and MDM2 isoforms showed that the N-terminal truncated short isoform of these proteins is less stable than the long isoform, suggesting that AS in CDS not only affects the functional domains but also changes the protein stability. We next asked whether mTOR-driven AS in CDS also affects proteome-wide post-translational modifications (PTMs) regulated by mTOR signaling^30–33^. We catalogued unique peptide sequences belonging to each AS isoform and compared them to peptide sequences in published PTM proteomic. In this proteome-wide search, we found that exon skipping could abolish existing sites for four interrogated PTMs in select genes and could also create new PTM sites in other genes, demonstrating that mTOR-modulated AS serves as a molecular scaffold for PTMs in functional proteomics (Fig. 3C). For example, the long SIRT2 isoform (NM_022432) creates a unique phosphoserine site that allows for cell cycle-dependent chromatin localization while the short isoform constitutively localizes to the cytoplasm^34,35^.

**Figure 3.**
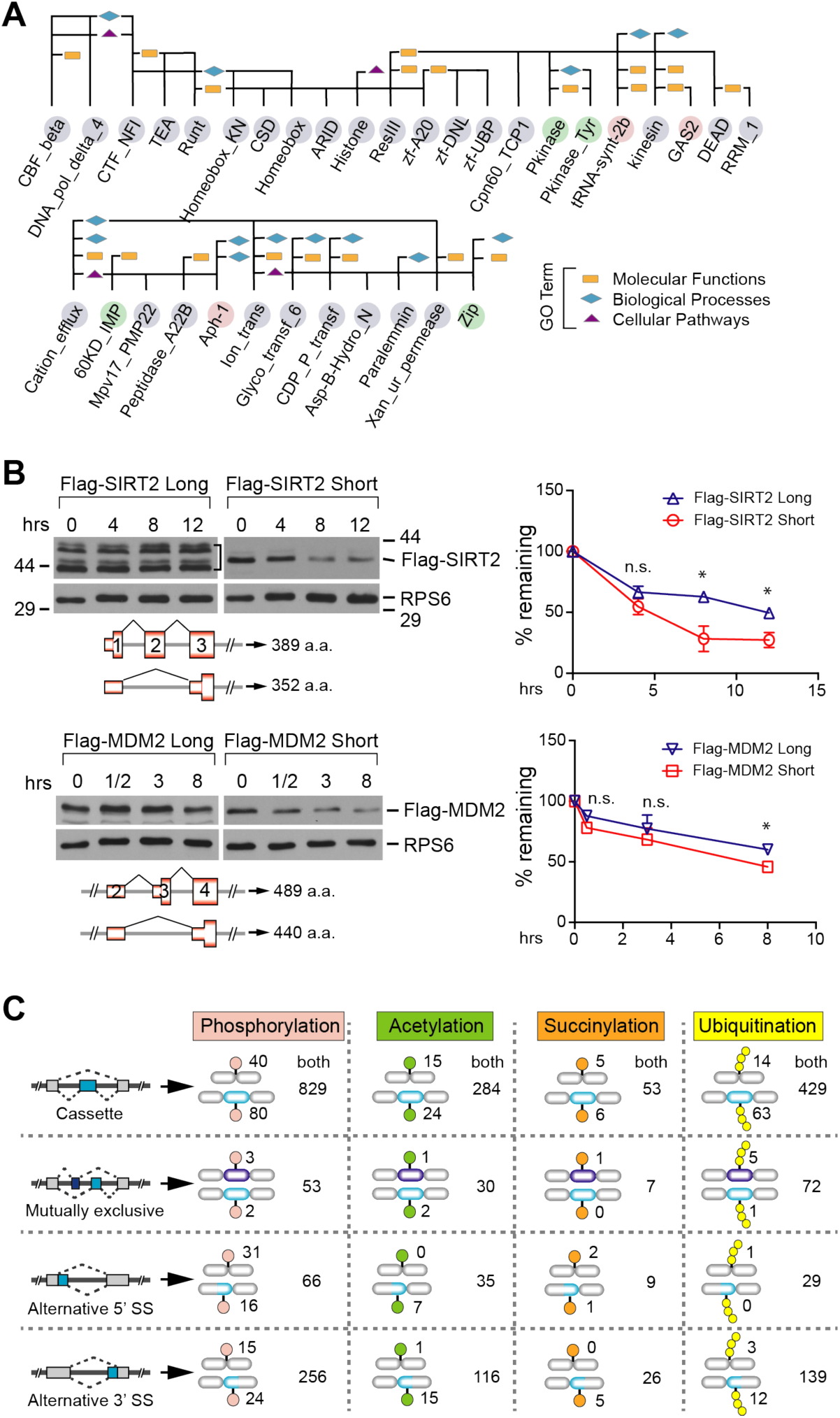
mTOR-driven exon skipping in CDS confers multifarious features in the proteome. A) Examples of affected Pfam domains and their associated GO terms by mTOR-driven AS events. Bioinformatics analysis of the functional Pfam domains disrupted or gained by AS events in the CDS of mRNA was performed. Pink: Pfam domains gained in hyper-activated mTOR. Light purple: Pfam domains lost in hyper-activated mTOR. Green: Pfam domains affected in both low and high mTOR environments. Rectangle: Molecular Functions; Diamond: Biological Processes; Triangle: Cellular Pathways. B) Analysis of protein isoform stability by western blotting. Flag-tagged SIRT2 and MDM2 protein isoforms were transiently expressed in HEK293 and the difference in their stabilities was monitored in the presence of cycloheximide (30 ug/ml) for the indicated time points. Ribosomal protein S6 (RPS6) was loaded as a control. The protein level was quantified using densitometry in ImageStudioLite software and normalized to the S6 level. Please note that the long Flag-MDM2 isoform shows up as multiple bands in the western blot. Mean (SEM) from two technical repeats were subjected to two-tailed Student’s t-test for statistical analysis. P<0.05 as significant (*). n.s. denotes no significance. C) Various post-translational modification (PTM) sites found on protein isoforms generated by mTOR-regulated AS. Protein isoforms produced by four types of AS are presented from top to bottom. Four types of PTM (phosphorylation, acetylation, succinylation, and ubiquitination) were interrogated using published mass spectrometry datasets. The number of unique PTM sites found in either exon-included or exon-skipped protein isoform is summed. The number of common PTM sites located in both protein isoforms is also reported.

### SRSF3 promotes exon skipping in the mTOR-activated transcriptome

Previously we reported the upregulation of splicing factor SRSF3 by mTOR-driven 3’-UTR APA and suggested a suppressive role of SRSF3 in mutually exclusive AS^36^. To more broadly understand the effect of mTOR-activated SRSF3 upregulation on transcriptome-wide AS, we knocked down *Srsf3* expression in *Tsc1*^-/-^ MEFs and profiled AS events by RNA-Seq experiments (Fig. 4A). Interestingly, we found that 60% of cassette exon-inclusion cases are preferred in the *Srsf3* knockdown cells, suggesting that SRSF3 promotes exon-skipping. Further analysis of overlapping exon skipping events by SRSF3 and mTOR activation in *Tsc1*^*-/-*^ MEFs showed that there is a 12.9% (64/497 cases) overlap (Fig. 4B), suggesting that SRSF3 upregulation is responsible for mTOR-dependent exon skipping. A similar observation was made when the *Srsf3* knockdown data were compared with the dataset of WT versus *Tsc1*^-/-^ MEFs (16.1%) (Suppl Fig. 4A). This suppressive role of SRSF3 on select genes was validated using RT-PCR (Suppl Fig. 4B). To gain an insight into the suppression mechanism, an *in silico* search was conducted to map the consensus binding sequence of SRSF3 across the suppressed exon regions. This approach identified two major patterns of SRSF3-binding: a cluster of SRSF3 consensus motifs located immediately upstream of the 3’-splice site and nearly the entire suppressed exon regions (Fig. 4C). Previous transcriptome-wide footprinting for SRSF3 showed that SRSF3 binds to exon regions and prefers penta-pyrimidine sequences^37^. Thus, mTOR activation-dependent SRSF3 upregulation could suppress exon inclusion by competing for polypyrimidine tracts located upstream of the 3’-splice site. It could also compete with splicing enhancers to bind exons and suppress exon inclusion in AS.

**Figure 4.**
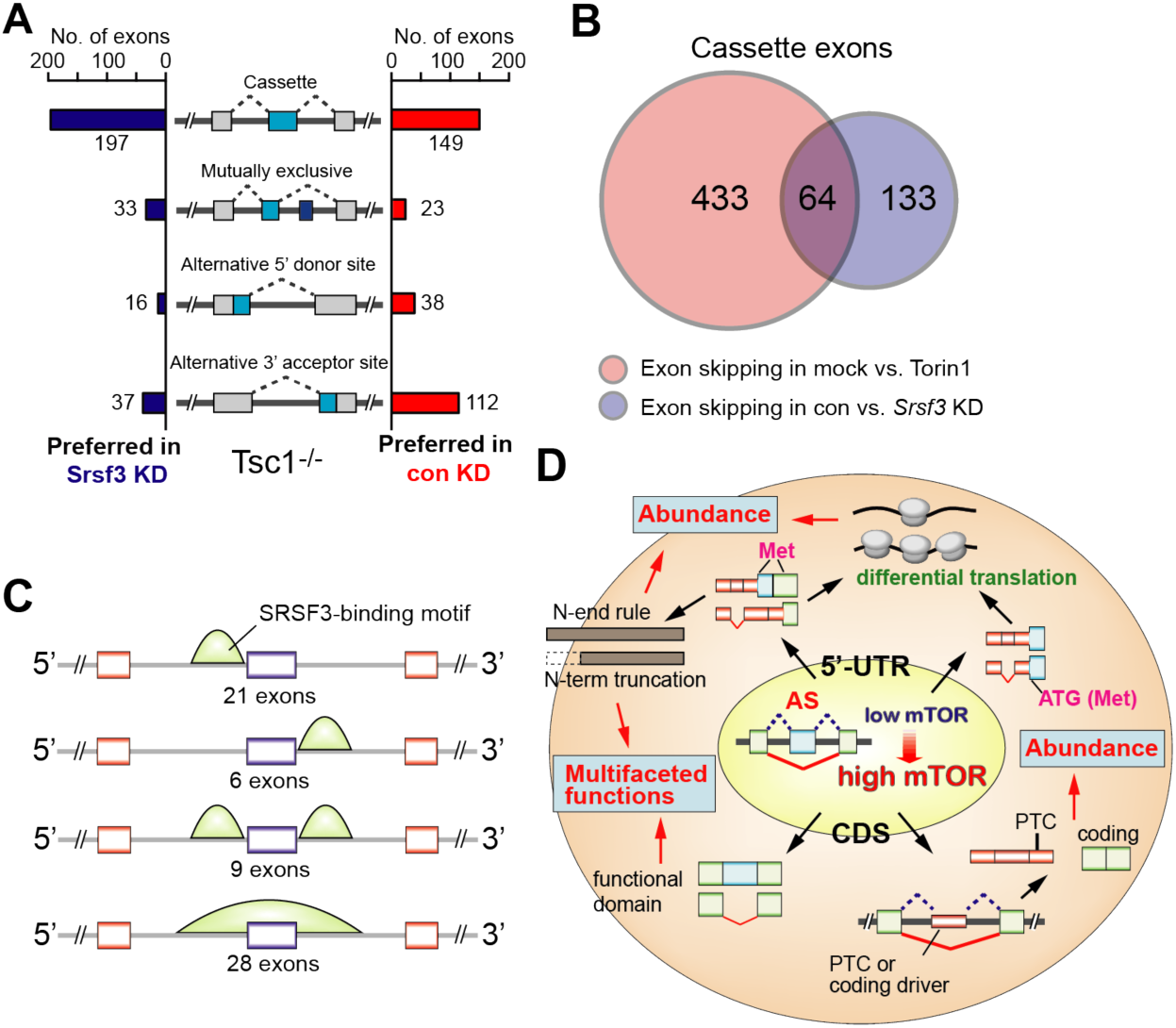
SRSF3 suppresses exon inclusion in mTOR-activated cells. A) Analyses of four different types of alternative splicing events in control vs. *Srsf3* knockdown in *Tsc1*^-/-^ MEFs. Cases of alternative exon inclusion are compared and illustrated. B) Venn diagram illustrating the overlap of skipped exons in mock vs. Torin 1 in *Tsc1*^-/-^ MEFs and control vs. *Srsf3* knockdown in *Tsc1*^-/-^ MEFs datasets. C) *In silico* prediction of SRSF3 putative binding sites using the overlapped exons as reported in B). The putative sites were determined using RBPMap^38^. D) Summary model of mTOR-mediated AS and the impact on the functional proteome.

This study identified mTOR-promoted transcriptome-wide exon skipping. These global AS events regulate the proteome multifariously: AS in CDS renders functional diversity and abundance of protein while AS in 5’-UTR mostly controls protein abundance (Fig. 4D). Further studies will be needed to understand the physiological relevance of AS in mTOR biology. Our study also identified SRSF3 as a key RNA-processing protein involved in mTOR-activated widespread exon skipping. Additional mechanistic experiments will be needed to further delineate other RBPs responsible for mTOR-regulated AS.

## Acknowledgements

We would like to thank Dr. Kwiatkowski at Harvard University for providing us *Tsc1*^*-/-*^ MEFs and matching wild type MEFs. We would also like to thank Dr. Yue Chen for providing us the lysine acetylation and succinylation proteomics datasets. We would also like to acknowledge Dr. Ann Hertzel for critically reading the manuscript and providing feedback.

## Declarations

The authors declare no conflict of interest.

## Methods

### Cell Lines

WT, Tsc1-/- MEFs, HEK293, MCF7, and MDA-MB-361 cells were cultured in Dulbecco’s Modified Eagle Media (DMEM) (Gibco), in supplements with 10% FBS (Gibco), 100ug/ml streptomycin, and 100U/ml penicillin. All cell lines were cultured at 37°C with 5% CO2.

### Plasmids

cDNAs of mouse Sirt2 long form (NM_022432), Sirt2 short form (NM_001122765), mouse mdm2 long form (NM_010786.4), mdm2 short form (NM_001288586.2) were cloned into p3xFlag-CMV-14 plasmid.

### Real-time PCR (RT-PCR) and Real-time quantitative PCR (RT-PCR) analysis

Total RNAs were isolated using the Trizol method recommended by manufacturer’s protocol. cDNAs were made by reverse transcription using Oligo-d(T) and superscript III (Thermo Fisher Scientific). SYBR Green fluorescence was used for quantitative real-time PCR reactions. The relative expression between groups was measured using the ΔΔCt method. The list of primers used for RT-PCR and qRT-PCR is:

mouse *Camkk2* RT-PCR forward: 5’-GGGAACCCGTTCGAAGGTAG-3’; mouse *Camkk2* RT-PCR reverse: 5’-GCAGGGACCACCTTTCACAA-3’; mouse *Ranbp3* RT-PCR forward: 5’-AAGCCTGCCGTCGCACCGTCTGTCT; mouse *Ranbp3* RT-PCR reverse: 5’-CTTCTCCGGCTTCGGGACTGGAGC-3’; mouse *Mdm4* RT-PCR forward: 5’-GCTAAGAAAGAATCTTGTTACATCAGC-3’; mouse *Mdm4* RT-PCR reverse: 5’-ATGTCGTGAGGTAGGCAGTGTGTGA-3’; mouse *Syce2* RT-PCR forward: 5’-AGCATCGGCAGAGTGAGAAC-3’; mouse *Syce2* RT-PCR reverse: 5’-CCGTTTCCACAGTTTGGCAG-3’; mouse *Itch* RT-PCR forward: 5’-TTCACAGTGGCCTTCTGGAAACAACG-3’; mouse *Itch* RT-PCR reverse: 5’-GGAATCAAGCTGTGGTCCACTGTCAGA-3’; mouse *Cenpi* RT-PCR forward: 5’-AGAGGCTGTGTGCTCGCCGAGGT-3’; mouse *Cenpi* RT-PCR reverse: 5’-TGAAAGCAGATGACTGGCCTACAGTAG-3’; mouse *Gabpb2* forward: 5’-GAGAACCAACTCCTGCGAGTTGTC-3’; mouse *Gabpb2* reverse: 5’-TCCCCAAGTCCACCAGAGACATCTTA-3’; human MDM4 forward: 5’-CGTCAGAGCTTCTCCGTGAAAGAC-3’; human MDM4 reverse: 5’-CTCTAGAATGTATGCATTTATGCTC-3’; mouse *Eef2* RT-PCR forward: 5’-AGTGTCCTGAGCAAGTGGTG-3’; mouse *Eef2* RT-PCR reverse: 5’-AGATCAGCGGTGAAGCCAAA-3’; mouse *Hsp90ab1* RT-PCR forward: 5’-CAGAAGGCTGAGGCAGACAA-3’; mouse *Hsp90ab1* RT-PCR reverse: 5’-AATCATGCGGTAGATGCGGT-3’; mouse *Anks1* RT-PCR forward: 5’-GCTGACTCGAAAGGCTGCTACC-3’; mouse *Anks1* RT-PCR reverse: 5’-TCCAGGGGCGTTTCAAACTTGTTG-3’; mouse *Ect2* RT-PCR forward: 5’-GAAATGCCGCAGGTTGAAGCAAG-3’; mouse *Ect2* RT-PCR reverse: 5’ ACTGGTGGCCCAACAATCCTACA-3’; mouse *Adgrl2* RT-PCR forward: 5’-TCCTCTGTGAGGCTGATGGAAC-3’; mouse *Adgrl2* RT-PCR reverse 5’-CAACAACATTGTGGCTGTGTGCG-3’.

### siRNA knockdown and antibodies

Cells were transfected with siRNAs synthesized by Integrated DNA Technologies (IDT) using RNAiMax Transfection reagents (Thermo Fisher Scientific) recommended by the manufacturer’s protocol. siRNA targeting mouse *Srsf3*: 5’-CGUGAUAUCAAGAAUUGU-3’. Antibodies used for western blot analysis include: anti-Flag (No. F3165, Sigma Aldrich), anti-S6 (No. 2317, Cell Signaling), Secondary antibody against mouse: goat-anti-mouse IgG-HRP (No. sc-2005, Santa Cruz Biotechnology), Secondary antibody against rabbit: goat-anti-rabbit IgG-HRP (No. sc-2004, Santa Cruz Biotechnology).

### Polysome isolation and analysis

Polysomes from mock-treated and Torin1-treated Tsc1^-/-^ MEF cell lysates were isolated using a sucrose gradient^40^. Cells were lysed in lysis buffer containing 20mM Tris pH=7.4, 150mM NaCl, 5mM MgCl_2_, 1mM dithiothreitol, 100mb/ml cycloheximide, 1% TritonX-100. Cytoplasmic extracts were loaded onto the sucrose gradient of 5% to 45% followed by ultracentrifugation of 36,000 rpm for 2 hours at 4°C. Eleven fractions were collected in total. Total RNAs were isolated using the Trizol method stated above. Ten percent (v/v) of the total RNAs from each fraction was used for cDNA synthesis for RT-PCR analysis. The total intensity was determined from the band intensity of the transcript isoform of each fraction combined and set as 100%. The relative ratio of the band intensity from each fraction over the total intensity was calculated and was used to generate the cumulative plots. Data presented shows the comparison of the two AS transcript in each condition (mock and Torin 1) For each cumulative plot, the x-axis denotes the fraction number, whereas the y-axis is the cumulative probability. To check the statistical difference between the two AS transcript isoforms in each condition, the Kolmogorov-Smirnov^39^ was applied and the *p-*value was reported in each plot.

### mRNA stability

Cells were treated with 1ug/ml actinomycin D to inhibit transcription and were harvested at different time points in the course of 12 hours. Total RNAs were obtained using the Trizol method stated above.

### Protein stability

Cells were treated with 30ug/ml cycloheximide to inhibit protein synthesis and were harvested at different time points. Cells were then lysed and lysates were run on SDS PAGE gels for western blot analysis.

### RNA-seq Data Analyses

To evaluate the transcriptome features under mTOR-hyperactivation at single-nucleotide resolution, we performed RNA-Seq analyses of poly(A+) RNAs isolated from WT, *Tsc1*^-/-^ MEFs, MCF7 and MDA-MB-361 cells. Paired-end reads were aligned to the mouse mm10 reference genome or human hg19 reference genome using TopHat2^40^ with up to two mismatches allowed. Kallisto^41^ was applied to quantify gene expressions with RefSeq annotation^42^.

### AS-Quant (*A*lternative *S*plicing *Quant*itation) pipeline

Alternative splicing analysis was performed using the AS-Quant pipeline^22^. Briefly, the pipeline categorizes the alternative splicing events into cassette exon, mutually exclusive, alternative 5’ splice site, and alternative 3’ splice site. Reads (n) from these alternatively spliced exons were measured and compared to reads (N) coverage of the rest of the transcripts. A 2×2 Chi-Square test was performed to determine the statistical significance for each alternative splicing event using the ratio of n/N. Events with a p-value <0.1 and a ratio difference >0.1 were considered real alternative splicing exons.

### Pfam domain analysis

Pfam-Scan was used to search Pfam databases for the matched Pfam domains on each transcript with 1e-5 e-value cutoff^43^. Only the Pfam domains which were overlapped with skipped exon(s) were considered for the further analysis. The InterPro Protein Families Database^44^ provided the information to link the Pfam domains to the Gene Ontology terms. The LinkDB under the GenomeNet Database^45^ provided the information to link the Pfam domains to the KEGG pathways.

### Quantification and Statistical Analysis

Signal intensity from gel electrophoresis or western blots was quantified using the ImageStudio software. Statistical analysis was performed by Two-tailed Student’s test unless stated otherwise.

### Data Availability

The accession numbers for the RNA-seq data are SRP056624.

## Supplemental Figures

**Supplemental Fig. 1.**
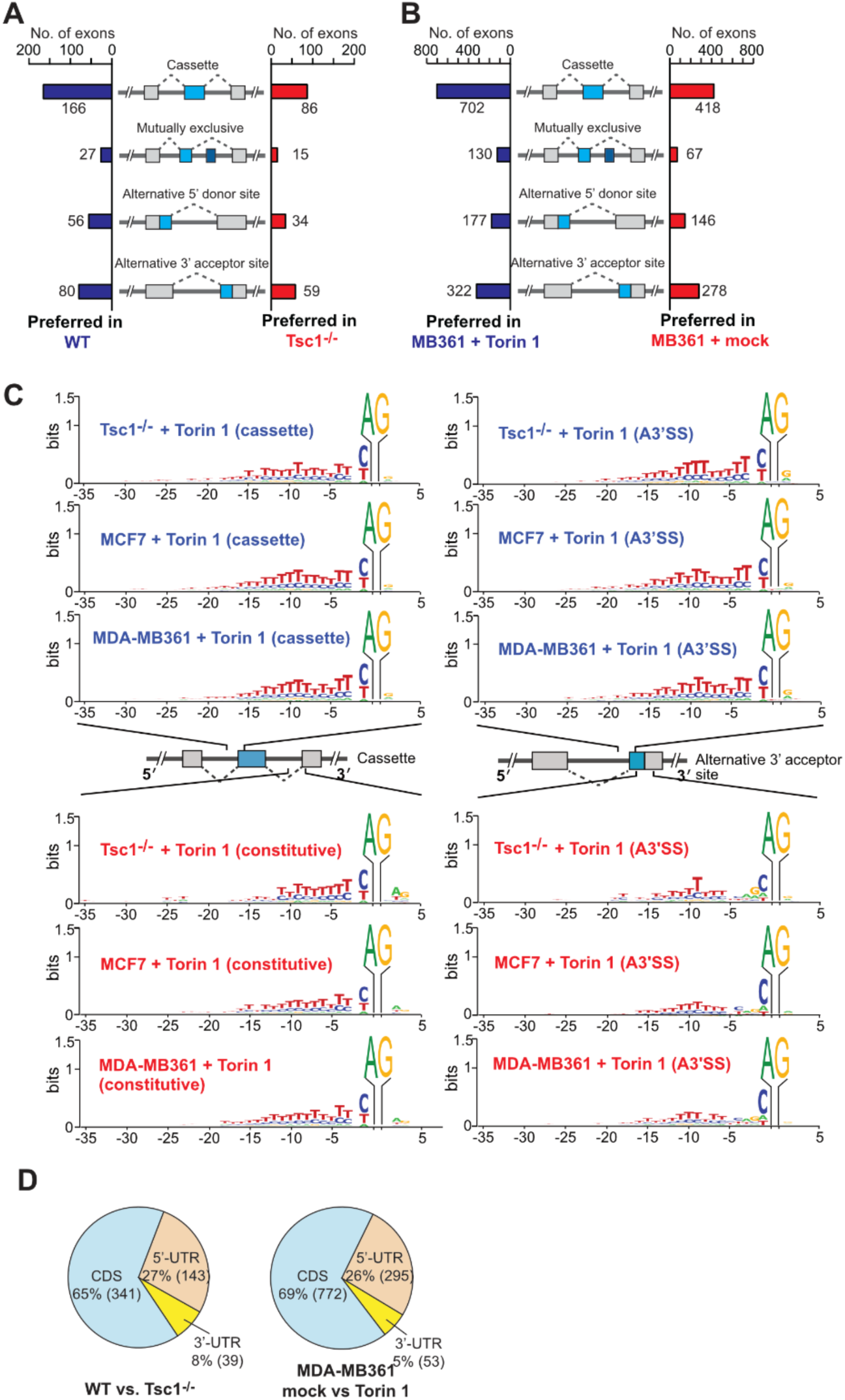
Cases of alternative exon inclusion when comparing A) WT vs. Tsc1^-/-^ MEFs and B) MDA-MB-361 mock vs Torin 1 treatment. C) The frequency of sequence preference of the 3’-splice site at AS exon compared to constitutively spliced exons for Tsc1^-/-^, MCF7, and MDA-MB-361 mock- and Torin 1-treated cells. X-axis represents the nucleotide position relative to the 3’ AG dinucleotide and Y-axis represents the certainty of the nucleotide. D) Distribution of mTOR-regulated alternative splicing (AS) events in different regions of the mRNA.

**Supplemental Fig. 2.**
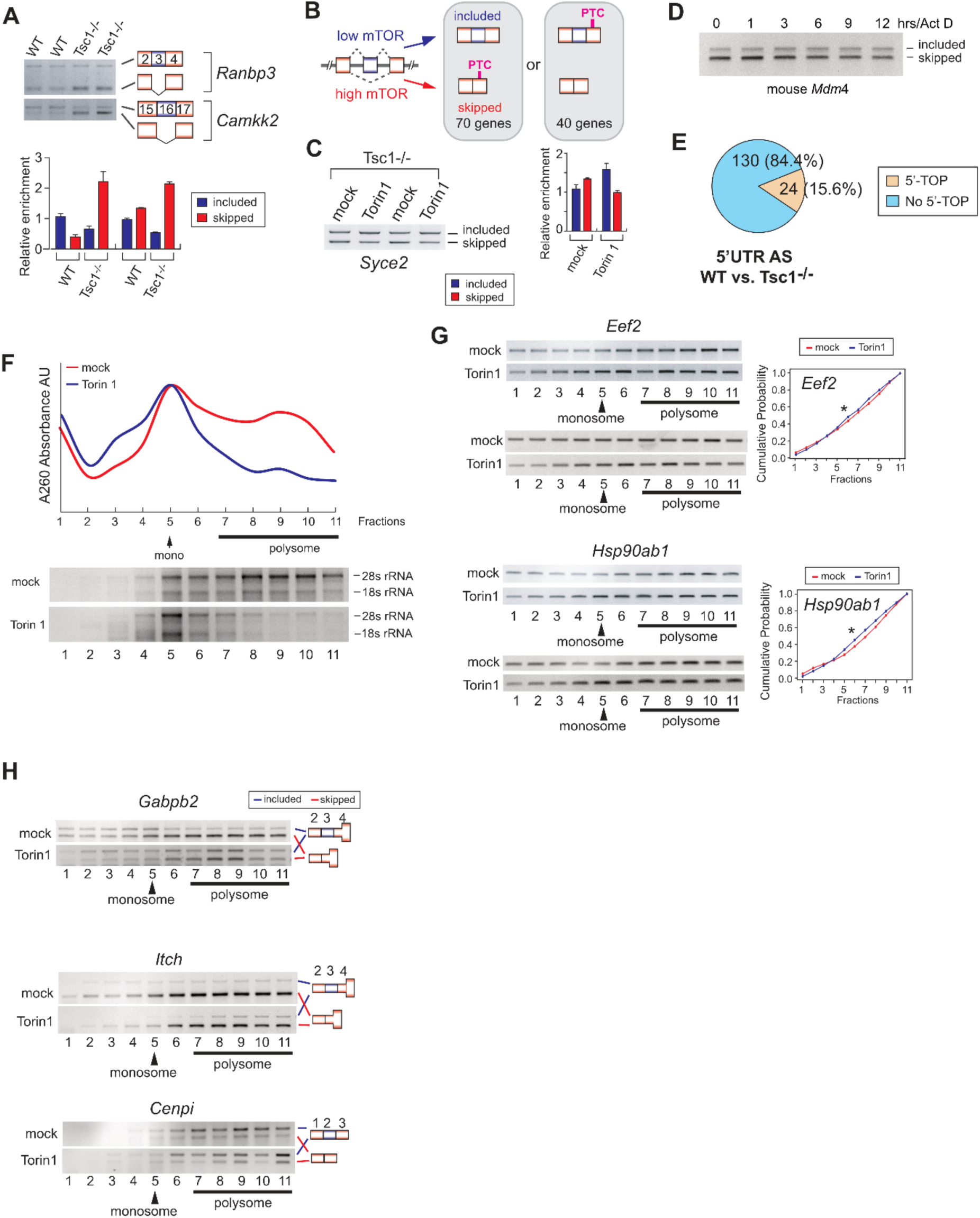
A) Semi-quantitative RT-PCR validation of mTOR-regulated AS events. (top panel) RT-PCR and analytical gel electrophoresis were conducted in WT and Tsc1^-/-^ MEFs. Two technical repeats were done. (bottom panel) Quantitation of gel images was done using ImageStudioLite. B) Schematic diagram illustrating the transition between coding and noncoding transcript caused by AS of cryptic exons. The number of genes showing coding to noncoding or vice versa by mTOR-driven AS events is shown. C) Semi-quantitative RT-PCR validation of coding-to-noncoding isoform switches. *Syce2* AS event was analyzed in Tsc1^-/-^ MEFs in the presence or absence of Torin1 treatment (50 uM). The relative quantification of the coding and noncoding transcripts was done using densitometry analysis. D) Technical repeat of stability of *mdm4* isoform transcripts. Semi-quantitative RT-PCR analysis was performed using Tsc1^-/-^ MEFs treated with 1ug/ml actinomycin D for indicated time points. E) Proportion of 5’-UTR AS events containing the 5’TOP motif in WT vs Tsc1^-/-^ MEFs. F) A260 absorbance plot and denatured RNA agarose electrophoresis of total RNAs from polysome profiling sucrose fractions in mock- and Torin 1-treated Tsc1^-/-^ MEFs. G) Polysome analyses of 5’-TOP containing transcripts *eef2* and *hsp90ab1.* Semi-quantitative RT-PCR and analytical gel electrophoresis were performed on the cytoplasmic fractions of mock- and Torin 1-treated Tsc1^-/-^ MEFs. The monosome and polysome fractions are indicated. The cumulative distribution of the transcript in each fraction is plotted against the summation of that isoform in all 11 fractions. The x-axis indicates the fraction number and y-axis indicates the cumulative probability. H) Technical repeat of polysome analyses of mTOR-regulated 5’-UTR AS transcripts.

**Supplemental Fig. 3.**
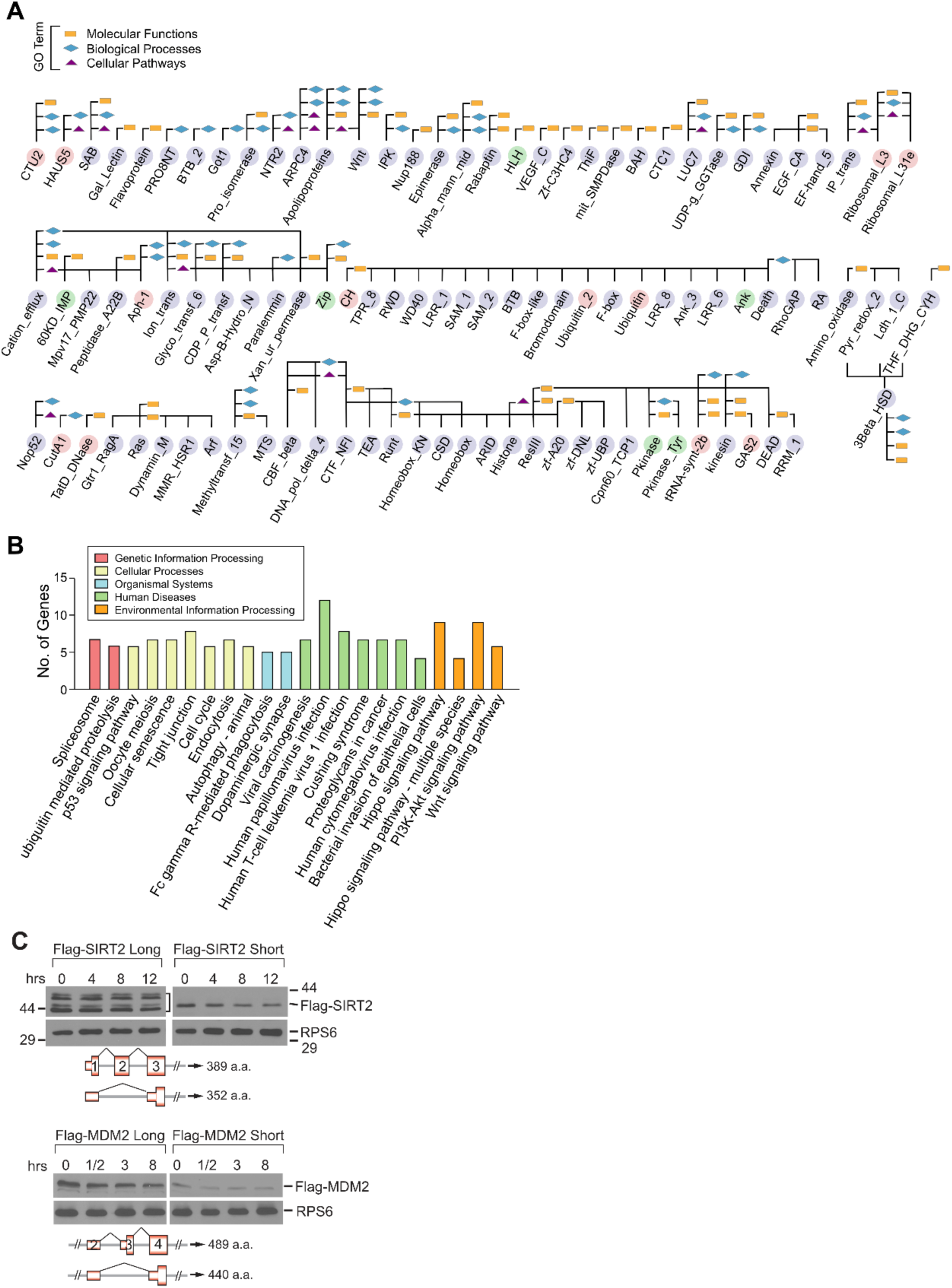
A) All of the impacted Pfam domains and their associated GO terms by mTOR-driven AS events. Bioinformatics analysis of the functional Pfam domains disrupted by AS events in the coding region of mRNA was done. Pink: Pfam domains gained in hyper-activated mTOR. Light purple: Pfam domains lost in hyper-activated mTOR. Green: Pfam domains present in both low and high mTOR environments. Rectangle: Molecular Functions; Diamond: Biological Processes; Triangle: Cellular Pathways. Each of affected Pfam domains associated with a GO term is indicated by a linkage to rectangle, diamond, and/or triangle. B) Affected KEGG pathways by mTOR-driven AS events determined using g:Profiler^44^. The KEGG pathways are grouped into 5 categories, shown by the legend. C) Technical repeat of protein isoform stability by western blotting. Flag-tagged SIRT2 and MDM2 protein isoforms were transiently expressed in HEK293 and the differences in their stabilities was monitored in the presence of cycloheximide (30 ug/ml) for indicated time points. Ribosomal protein S6 (RPS6) was loaded as a control. The protein level was quantified using densitometry and normalized to the S6 level.

**Supplemental Figure 4.**
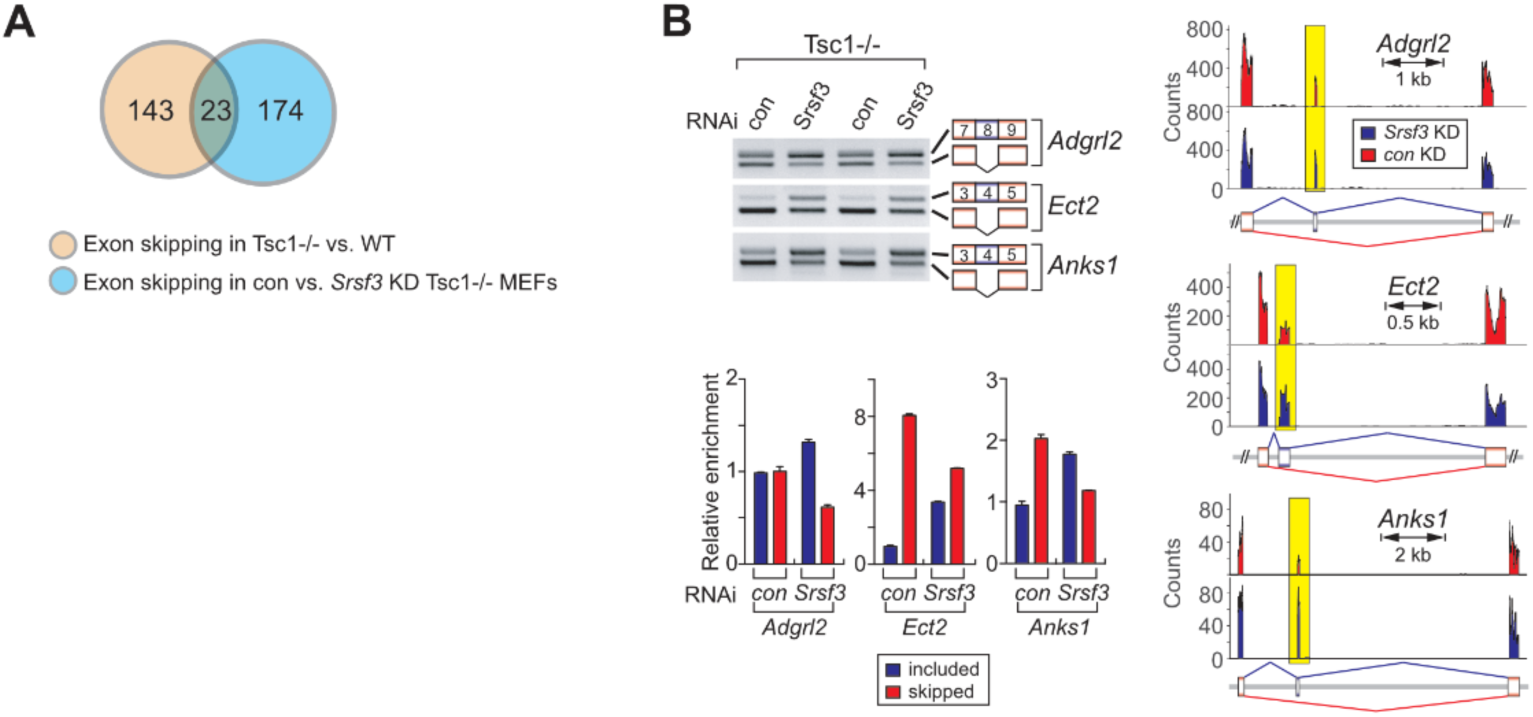
A) Venn diagram illustrating the overlap of skipped exons in WT vs. Tsc1^-/-^ MEF dataset and control vs. *Srsf3* knockdown in Tsc1^-/-^ MEFs dataset. B) Semi-quantitative RT-PCR validation of *Srsf*3 knockdown RNA-seq experiment. Alternative splicing of *Anks*1, *Ect*2, and *Adgrl*2 were tested in control knockdown and *Srsf*3 knockdown Tsc1^-/-^ MEFs. Tsc1^-/-^ MEFs were transfected with scrambled or *Srsf3* siRNAs for 48hrs, and total RNAs were harvested. mRNAs were reverse transcribed into cDNA, which served as a template for RT-PCR.

